# Spatial Rewiring of Enterocyte Identity in Celiac Disease

**DOI:** 10.64898/2026.02.14.705877

**Authors:** Tal Barkai, Rachel Frieman-Sharabi, Keren Bahar Halpern, Roy Novoselsky, Yael Korem Kohanim, Sapir Shir, Ofra Golani, Inna Goliand, Yoseph Addadi, Merav Kedmi, Hadas Keren-Shaul, Lena Prichislov, Anat Guz-Mark, Hadassah Nissim, Raanan Shamir, Dror Shouval, Shalev Itzkovitz

## Abstract

Enterocytes in the human small intestine exhibit distinct functional states in different zones along the crypt-villus axis, a feature that is thought to convey optimal absorption. In celiac disease (CeD), autoimmune destruction of enterocytes leads to villus blunting, but how this altered tissue morphology affects enterocyte states is unclear. Using spatial and single-cell transcriptomics, we show that in patients with CeD, enterocytes acquire a novel identity characterized by co-expression of multiple zonal programs. This aberrant zonal co-expression results from reduced distances between BMP- and WNT-producing mesenchymal cells, leading to overlapping morphogen fields. In addition, we identify a subset of metaplastic cells that adopt gastric pit cell-like identities in discrete tissue patches. Our findings provide a detailed view of epithelial remodeling in CeD and establish a resource for understanding the cellular basis of malabsorption associated with villus blunting.

## Introduction

The small intestine is composed of repeating crypt-villus units, facilitating the large surface area needed for absorption. Recent studies in mice and humans have shown that all intestinal cell types, including the nutrient absorbing enterocytes^1^, secretory cells^2^ mesenchymal and immune cells^3–5^ exhibit strikingly different cellular states, depending on their location along the crypt-villus axis. Zonation of enterocytes has been suggested to facilitate optimal absorption through division of labor^6^. In healthy humans, enterocytes at the base of the villi absorb micronutrients and transport antibodies into the lumen, whereas enterocytes at the middle and tip of the villi specialize in absorption of distinct macronutrients. Processes such as lipid absorption and iron processing are further distributed across multiple villi zones, facilitating temporal delays and tight homeostatic control^5^.

CeD is an autoimmune disorder that affects about 0.5-2% of the global population and occurs in genetically susceptible individuals^7–10^. In people with CeD, eating gluten triggers an abnormal immune response in which modified gluten fragments activate CD4⁺ T cells and stimulate the production of autoantibodies against gliadin and the enzyme tissue transglutaminase 2^11^. Innate immune pathways and inflammatory cytokines further amplify this response, driving intestinal damage^12^. This damage often leads to complete blunting of the intestinal villi, that leads to malabsorption and is associated with other symptoms, such as abdominal pain and diarrhea.

Although many of the immune mechanisms driving CeD have been characterized^13–17^, the consequences of villus blunting for epithelial cell identity and spatial organization remain poorly understood. Villus blunting could lead to selective loss of zone-specific enterocyte states, a proportional reduction across all zonal programs, or a complete breakdown of spatial zonation with intermixed enterocyte identities (Fig. 1a). Here, we address this question using single cell and high-resolution spatial transcriptomics. We uncover a strikingly different scenario, in which enterocytes are not simply lost but are rather reprogrammed to simultaneously express multiple zonal programs within individual cells. In addition, we identify extensive gastric metaplasia in discrete epithelial patches. These findings indicate that malabsorption in CeD arises not only from reduced epithelial surface area, but also from disruption of finely tuned zonal enterocyte states.

**Figure 1.**
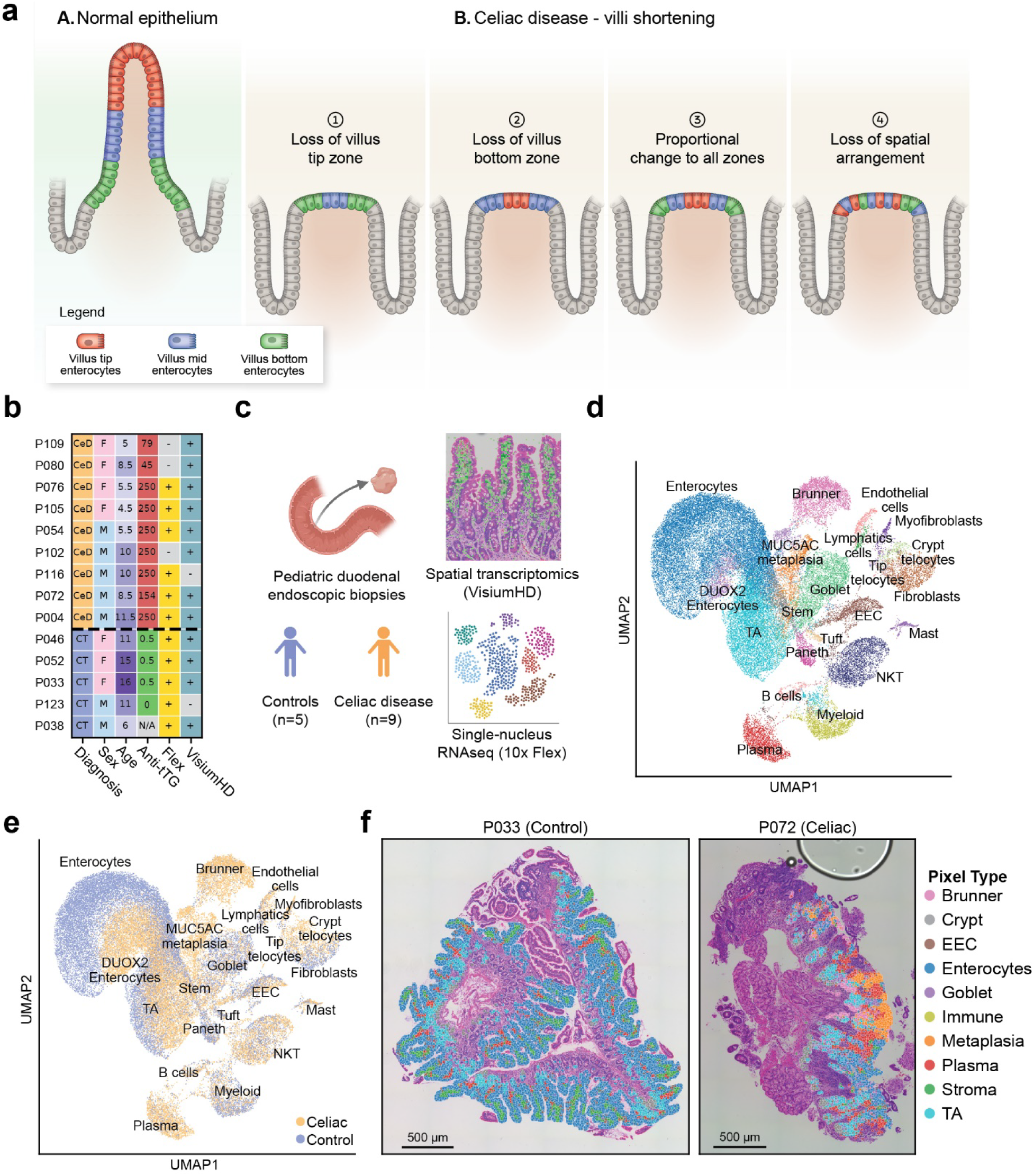
Spatially resolved single-cell profiling of pediatric duodenal biopsies in CeD and controls. **a.** (A) Normal villus zonation showing bottom, mid, and tip enterocytes. (B) Potential changes in blunted villi: loss of tip enterocytes (1), loss of bottom enterocytes (2), proportional loss across all zones (3), or disruption of spatial zonal organization (4). **b.** Patient characteristics and the technologies applied for each sample in our study. CT, control. Age in years, anti-tTG (tissue transglutaminase) serum levels are in U/ml. Anti-tTG values above 250 were fixed at 250. P038 is IgA-deficient, explaining the lack of anti-tTG values. **c.** Schematic overview of experimental design. Created using BioRender. **d.** Uniform Manifold Approximation and Projection (UMAP) projection of snRNAseq data annotated by cell type. **e.** UMAP projection of snRNAseq data annotated by condition. **f.** Representative spatial transcriptomics images from control (left) and celiac disease (right) samples, annotated by pixel type (Methods).

## Results

### Spatially resolved single cell atlas of CeD

To address the changes in zonal expression programs in CeD, we assembled a cohort of 9 pediatric patients referred for an esophagogastroduodenoscopy for suspected CeD. All 9 subjects were eventually diagnosed with CeD. In addition, 5 pediatric control subjects were analyzed. Patients in both groups were on a gluten-containing diet at the time of endoscopy. While all patients with CeD had typical histologic findings indicative of CeD (villous flattening, crypt hyperplasia and increased intra-epithelial lymphocytosis), all control subjects had normal endoscopic and histologic assessment of the duodenum (Fig. 1b). We acquired duodenal biopsies and applied VisiumHD spatial transcriptomics on H&E-stained sections, as well as 10X Flex single-nucleus RNAseq (snRNAseq) on dissociated tissues from sequential sections (Fig. 1c). Our snRNAseq atlas of 53,270 high-quality cells (Methods) included major immune, mesenchymal and epithelial cell types (Fig. 1d,e, Supplementary Fig. 1a-d). Similarly, our VisiumHD datasets included all cellular compartments (Fig. 1f, Supplementary Fig. 1e-h). We used the H&E images to classify each VisiumHD pixel into epithelium or stroma (Supplementary Fig. 1i, Methods). This combined dataset facilitated exploration of changes in zonated cellular states.

### Enterocytes in CeD undergo zonal reprogramming

To examine alterations in enterocyte zonation in CeD, we first used the VisiumHD datasets to reconstruct enterocyte organization along the crypt-villus axis in the control subjects. We annotated the crypt-villus boundary based on H&E images, and subdivided epithelial-specific pixels into consecutive 100 µm wide zones spanning the entire axis. For each zone, we calculated the average expression of all genes (Supplementary Fig. 2a-b, Supplementary Table 1, Methods). Using this zonal reconstruction, we identified enterocyte marker genes specific to the villus bottom, mid, and tip regions (Methods, Supplementary Table 2). We next examined these zonated markers in the epithelium of controls and of CeD patients.

In control subjects, enterocytes exhibited well-defined spatial gradients of zonal marker expression, consistently observed in both the snRNAseq (Fig. 2a) and VisiumHD (Fig. 2b,c) datasets. These gradients included crypt-restricted expression of *OLFM4*, expression of *REG1A* in the crypt and lower villus zones, mid-villus enrichment of the luminal glucose transporter *SLC5A1*, and villus tip-specific expression of the water channel *AQP10*. In the VisiumHD data, zonal enterocyte markers such as the bottom-to-mid villus gene *SLC2A2* and the villus tip gene *AQP10* showed clear spatial segregation (Fig. 2c). In gene expression space, enterocytes resided on a curve indicative of the sequential changes of enterocyte cell states during their migration from the bottom to the top of the villi (Fig. 2a, d).

**Figure 2.**
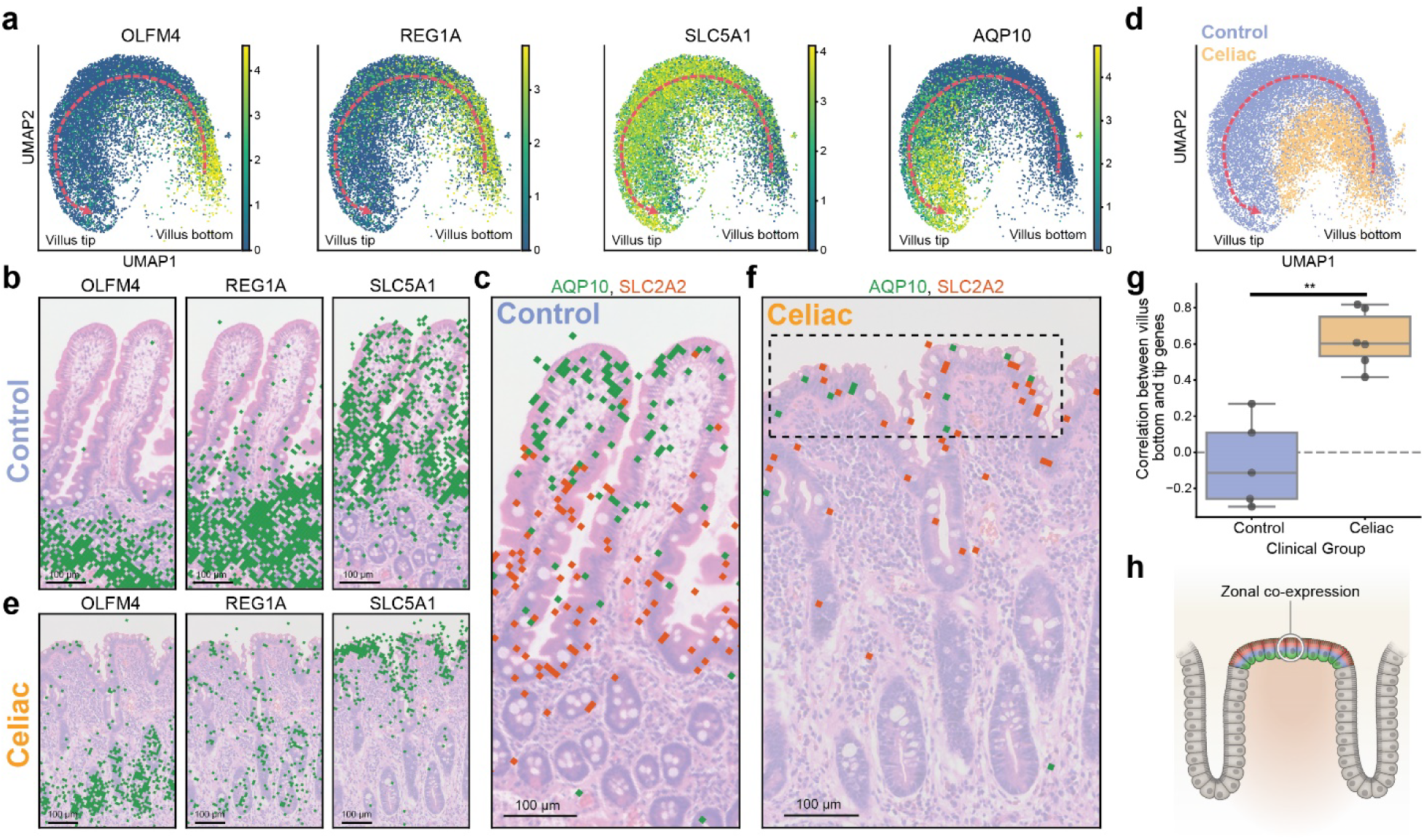
Enterocytes in celiac disease co-express multiple zonal programs. **a.** UMAP projections of enterocytes from control samples colored by the expression of zonal marker genes Expression scale – natural log of normalized counts (log1p). **b-c.** Spatial transcriptomics images from control sample showing expression of representative zonal enterocyte markers. **d.** UMAP projection of enterocytes annotated by disease condition. Dashed red curves in a, d are guides to the eye, indicating the direction of enterocyte migration from the villus base toward the villus tip. CeD enterocytes (orange) occupy a distinct gene expression sub-space compared to control enterocytes (blue). **e-f.** Spatial transcriptomics images from CeD sample showing the expression of the same representative zonal enterocyte markers of b. Dashed box in f highlights co-expression of genes that are zonally segregated in controls. **g.** Quantification of the Spearman correlation between bottom and tip zonal scores, demonstrating a significant difference in enterocyte zonal correlation of bottom and tip scores in CeD patients and controls. Each dot represents a patient; statistical significance was assessed using the Wilcoxon rank-sum test, *p*=0.006. Box plots show the median as the center line, boxes span the 25th–75th percentiles, whiskers extend up to 1.5 IQR. **h.** Scheme showing enterocytes with zonal co-expression programs in CeD.

In contrast to a scenario of selective loss of zonal enterocytes (Fig. 1a), CeD enterocytes did not span a subset of the control enterocyte zonated curve. Rather, CeD enterocytes occupied a gene expression space that was distinct from that of control enterocytes (Fig. 2d). Notably, enterocytes along the blunted villi in CeD co-expressed multiple zonal programs that were spatially segregated in controls (Fig. 2e,f). For example, *SLC2A2* and *AQP10* were expressed in distinct zonal enterocytes in control biopsies (Fig. 2c) yet were co-expressed within individual enterocytes in CeD biopsies (Fig. 2f, dashed box). To quantify this zonal co-expression, we used our snRNAseq atlas to compute single-cell correlations between villus bottom and villus tip marker genes in enterocytes (Fig. 2g, Supplementary Fig. 2c,d, Methods). These zonal programs were significantly anti-correlated in control patients (median R = −0.1, Fig. 2G, Supplementary Fig. 2c) but showed a significant positive correlation in CeD patients (median R = 0.6, Fig. 2g, Supplementary Fig. 2d) with significant difference between correlations of CeD patients and controls (*p*=0.006, Fig. 2g). Beyond the zonal reprogramming of CeD enterocytes, the global expression levels of villus enterocyte markers were lower in CeD, and the CeD intestinal crypt showed elevation of tip villus markers (Supplementary Fig. 2e,f). Together, these findings demonstrate that enterocytes in CeD are transcriptomically distinct from those in controls and exhibit aberrant co-expression of gene programs that are normally spatially segregated along the crypt-villus axis (Fig. 2h).

### Zonal reprogramming of enterocytes is associated with overlapping morphogen fields

What mechanisms could give rise to the zonal co-expression of enterocyte programs in CeD? Gene expression along the crypt-villus axis is shaped by morphogen fields emanating from mesenchymal cell populations^3,4,18–26^. As enterocytes migrate from the bottom of the crypt to the tip of the villus, they sequentially encounter decreasing levels of canonical WNT signals and increasing levels of BMP signals. BMP is critical for down-regulating bottom villus programs and up-regulating tip villus programs^22^. Our snRNAseq atlas included central mesenchymal cell types, notably *MYH11*+ myofibroblasts, *FBLN1*+ crypt telocytes and *NRG1*+ tip telocytes (Fig. 3a,b). These three cell types are the main sources of the key zonated WNT and BMP ligands (Methods, Supplementary Fig. 3a). Using our snRNAseq atlas, we found that the expression levels of the BMP and WNT markers in these mesenchymal cell types did not significantly differ between CeD and controls (Supplementary Fig. 3b,c).

**Figure 3.**
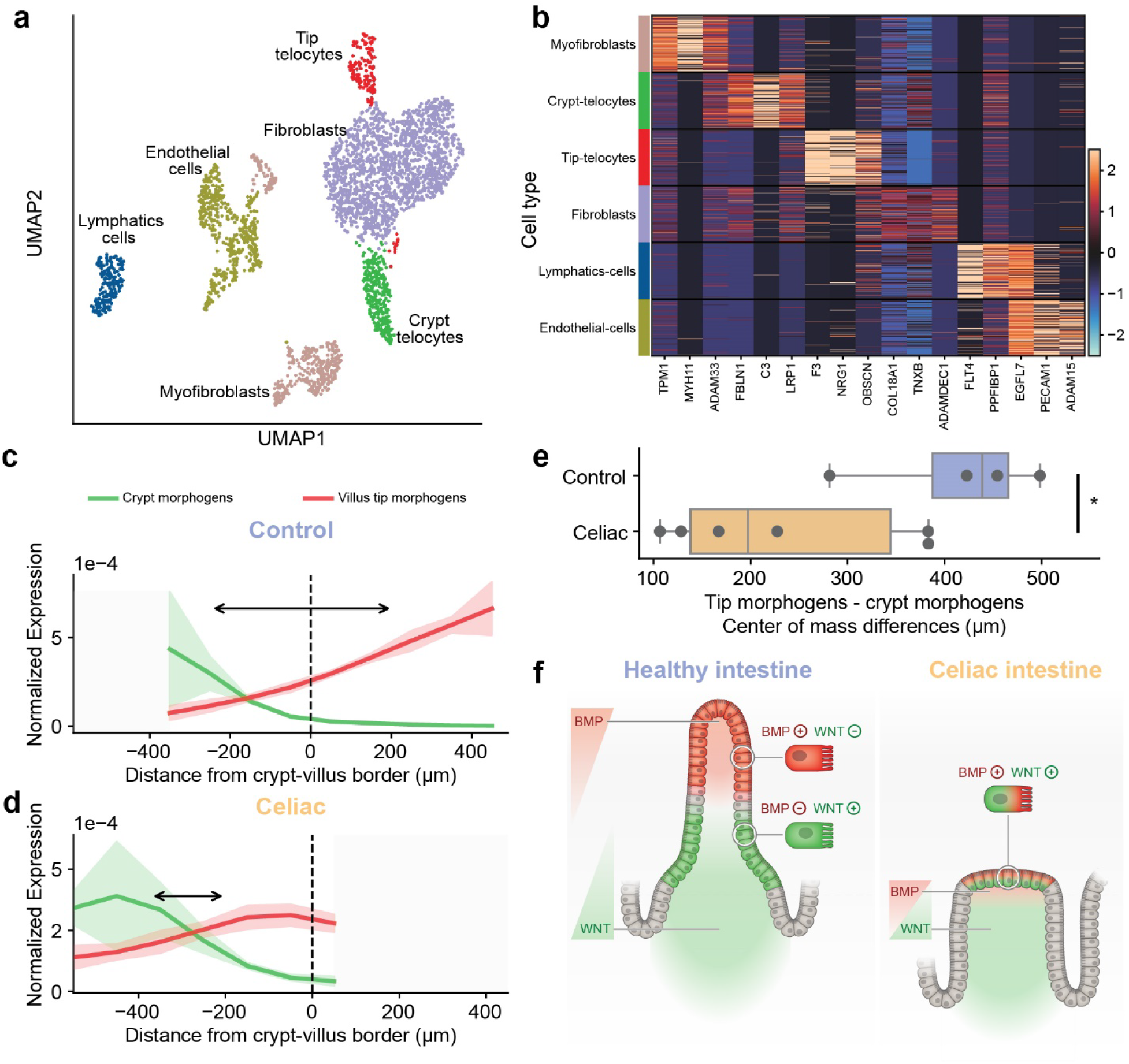
Villus blunting in CeD gives rise to overlap between BMP and WNT morphogen fields. **a.** UMAP projection of snRNAseq of the stromal compartment, clusters with less than 50 cells are not shown. **b.** Heatmap showing normalized expression of marker genes of each cell type cluster. Cells were sub-sampled to display 140 cells per each cluster. **c-d.** Sums of crypt (green) and tip (red) morphogen expression in celiac and control (Methods). Patches are standard errors of the means (SEM), profiles were smoothed with a Gaussian filter with σ=1. Arrows show differences between profiles’ COM. Gray zones are due to crypt hyperplasia and villus blunting. **e.** Patient specific crypt and tip COM differences. *p* = 0.033, rank-sum test. Box plots show the median as the center line, boxes span the 25th–75th percentiles, whiskers extend up to 1.5 IQR. **f.** Model of changes in morphogen gradients between normal and blunted villi. Morphogen fields are non-overlapping in controls and overlapping in CeD.

We used the VisiumHD datasets to compare zonal expression profiles of the BMP and WNT markers in the stromal pixels. We found that while the absolute expression levels of the crypt and tip morphogens were similar, the distance between their profiles’ centers of mass (COM, Methods) was significantly lower in patients with CeD vs. controls (median COM difference between the profiles’ COM of 197µm vs. 438µm, *p* = 0.033; Fig. 3c-e). This result suggests that the blunted villus morphology reduces the distance between the crypt and villus tip signaling sources, potentially creating overlapping rather than distinct morphogen fields. Migrating CeD enterocytes may therefore encounter mixed WNT and BMP signals at the blunted villi, yielding the coordinated elevation in the expression of genes that were normally distinct (Fig. 3f).

### A subset of blunted villi epithelial patches exhibits gastric metaplasia

Our analysis demonstrated that CeD enterocytes are not a subset of zonal enterocytes but rather express a distinct functional state that includes co-expression of programs that are normally spatially distinct. Notably, our VisiumHD data revealed patches of inter-crypt blunted villi with very low expression of enterocyte markers, regardless of zone (Fig. 4a). Rather, these patches exhibited elevated levels of gastric pit cell genes, including *MUC5AC*, *GKN1* and *GKN2* (Fig. 4a, Supplementary Fig. 4a-c)^27,28^. This metaplastic expression pattern resembled the recently reported surface foveolar cells observed in diverse intestinal pathologies^29–31^. *MUC5AC*+ cells formed a distinct cluster in our snRNAseq atlas (Fig. 4b), appeared in all CeD patients and formed a significant fraction of epithelial cells (7%±1.8% vs. 0.8%±0. 8%; Wilcoxon rank-sum test, *p* = 6.17×10⁻³). Comparison of the metaplastic cells to CeD enterocytes in our snRNAseq atlas revealed substantial gene expression differences, including a decrease in absorption programs and an increase in mucin biosynthesis programs (Fig. 4b-d, Supplementary Fig. 4d). Consistently, staining with wheat germ agglutinin, which binds mucus sugar residues, demonstrated a distinct mucus layer bordering the *MUC5AC*+ cells (Supplementary Fig. 4e). *MUC5AC* metaplastic cells further showed elevated expression of *FOXQ1*, a stomach-enriched transcription factor that serves as a key regulator and marker of differentiated gastric pit cells^32^. The blunted epithelium in CeD is therefore altered beyond zonal co-expression, with discrete cell patches fully losing enterocyte identity and adopting gastric pit cell-like states.

**Figure 4.**
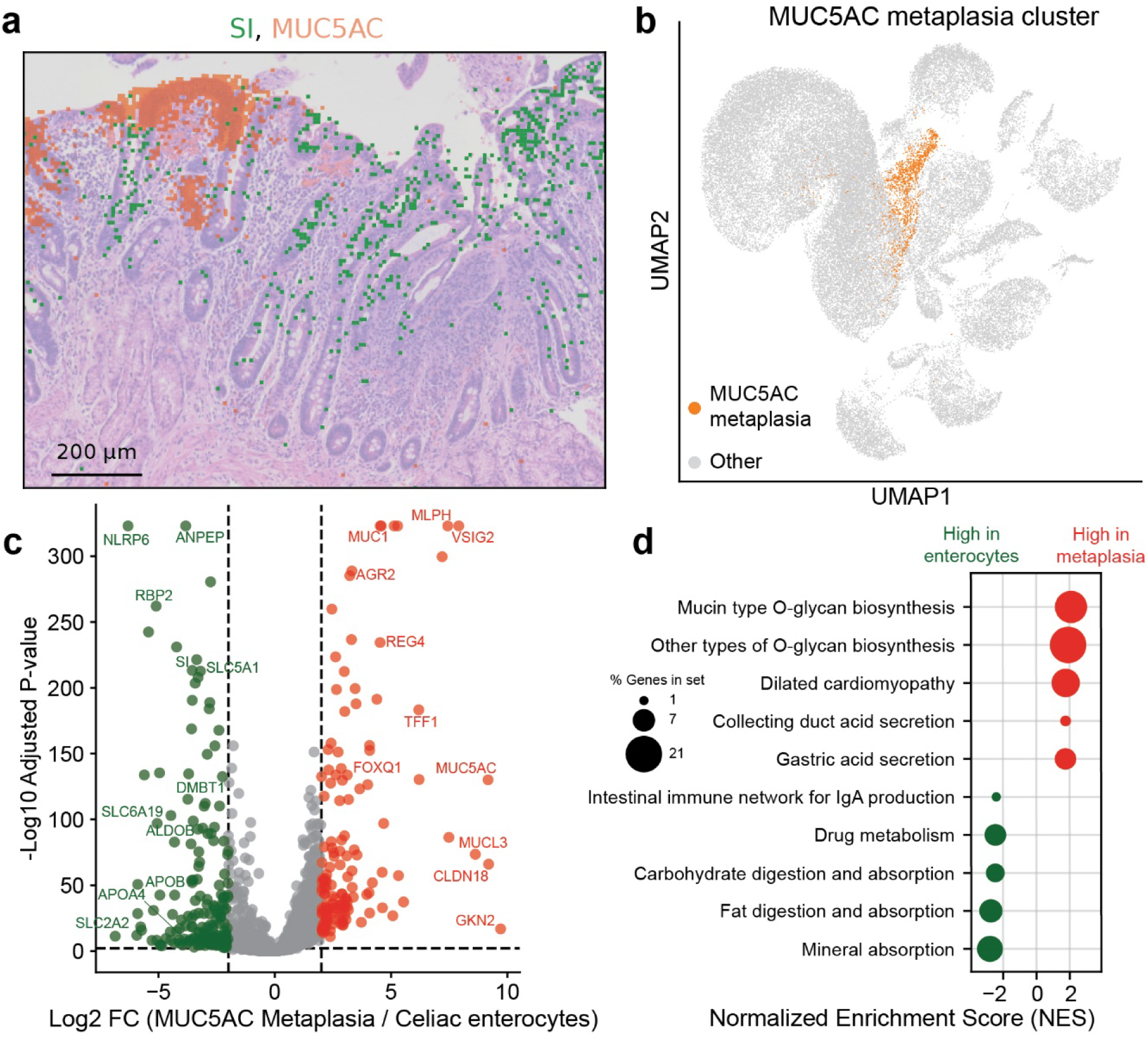
Gastric metaplasia in CeD **a.** VisiumHD image showing expression of an enterocyte marker gene Sucrose Isomerase (*SI*) and a gastric metaplasia marker gene (*MUC5AC*) in spatially distinct tissue regions. Image representative of 4 patients. **b.** UMAP projection of the snRNAseq data, highlighting the *MUC5AC*+ metaplastic cell cluster. **c.** Volcano plot showing differentially expressed genes between metaplastic *MUC5AC*+ cells and celiac enterocytes. Significance thresholds were set to a minimum absolute log₂ fold change of 2 and an adjusted *P* value ≤ 0.01. Shown are representative genes. **d.** Gene set enrichment analysis (GSEA) demonstrating enrichment of mucin biosynthesis pathways in metaplastic cells and classical enterocyte functional programs in celiac enterocytes.

## Discussion

CeD is associated with malabsorption of nutrients, leading to pathologies such as anemia, growth delays and osteopenia^7,9^. These effects have been attributed to the loss of small-intestinal surface area associated with the immune-mediated villus blunting. Here, we show that beyond the loss of enterocyte cell numbers, remaining enterocytes in CeD exhibit substantial changes in their functional cell states. These entail the co-expression of enterocyte programs that are normally distinctly expressed in different villi zones. The zonated gene expression patterns of enterocytes have been suggested to facilitate optimal absorption^5^. For example, the key steps of fatty acid absorption, entailing luminal fatty acid import, triglyceride biosynthesis, chylomicron assembly and release, are distributed across multiple villus zones, facilitating a delay in lipid release from enterocytes. This delay has been hypothesized to prevent spikes of blood lipid levels after meals^5^. Similarly, enterocyte iron absorption and release are zonally separated, presumably to enable tighter control of iron levels. Future metabolic studies could examine the physiological impact of the co-expression of these and other zonally distinct programs on dynamic blood metabolite concentrations following meals. The zonal co-expression states we identified in CeD could be explained by the overlap of morphogen fields stemming from the shorter distance between crypt and villus tip stromal signaling sources. It would be important to explore whether similar zonal co-expression of enterocytes occurs in non-CeD related villus blunting conditions^10^.

Beyond the zonal co-expression, CeD epithelium contained patches of metaplastic cells that resemble gastric pit cells, exhibiting a secretory rather than an absorptive phenotype. Our high-resolution spatial transcriptomics datasets demonstrated that, in CeD, metaplastic cells reside in blunted villi patches which lose expression of classic enterocyte absorptive genes. Intestinal gastric metaplasia has been described across diverse disease conditions in the human intestine^31,33–36^. Moreover, *MUC5AC*+ cells have been previously described in CeD^29–31^. The elevation in gastric-like mucus production could be a response to the thinner mucus layer, brought about by loss of goblet cell mass due to villus blunting. Conversely, it could emanate from aberrant differentiation processes in the crypt. Future studies, using either *in-vivo* approaches or organoids could address the mechanism leading to this metaplastic process^37^.

The intestinal epithelium has a remarkable regenerative capacity. Upon shifts to gluten free diet (GFD), villi regrow and the normal crypt-villus morphology reappears. It will be interesting to apply our approaches to study whether the healthy zonal enterocyte states rehabilitates upon the shift to GFD, or whether the enterocytes or the stem cells retain a memory of the reprogramming process that occurred during the active phase of the disease^38^. Similarly, it will be interesting to explore whether metaplastic enterocytes disappear after the shift to GFD. Our study highlights a distinct enterocyte cellular state of zonal co-expression, and provides a spatially resolved resource for exploring additional aspects of the cellular landscape aberrations in CeD.

## Acknowledgments

SI is supported by the Moross Integrated Cancer Center, the Helen and Martin Kimmel Award for Innovative Investigation, the Yad Abraham Research Center for Cancer Diagnostics and Therapy, the Israel Science Foundation MAVRI program grant no. 1482/25, the European Union (ERC, GI-DYNAMICS, 101198168), the Swiss Society Institute for Cancer Prevention Research, and a Weizmann-Schneider joint research grant and a research grant from the Ministry of Innovation, Science and Technology, Israel. Optical imaging data was acquired at the de Picciotto Cancer Cell Observatory In memory of Wolfgang and Ruth Lesser of the Moross Integrated Cancer Center in the department of Life Science Core Facilities, Weizmann institute of science.

## Author contributions

TB contributed to study conception and design, data collection, experimental procedures, data analysis, and manuscript writing. RFS, KBH, and RN contributed to data collection and experimental procedures. YKK and SS contributed to data analysis. OG, IG, and YA contributed to the microscopy and image analysis. MK, HKS, and LP contributed to experimental procedures. AGM and HN contributed to patient recruitment. RS and DS contributed to study conception, patient recruitment, and manuscript writing. SI contributed to study conception, data analysis, and manuscript writing.

## Data and code availability

Pediatric crypt-villus gene expression profiles in controls and CeD, as well as spatial transcriptomics images, can be browsed using our web app (https://shalevapps.weizmann.ac.il/celiac_atlas/).

The data and the computer code produced in this study are available in the following databases: Source code is available at GitHub (https://github.com/barkait/celiac_atlas), count matrices and background subtracted data of the Flex snRNAseq, microscopy images and count matrices of the VisiumHD datasets are available at Zenodo.

## Methods

### Sample collection and formalin-fixed, paraffin-embedded (FFPE) blocks preparation

Pediatric patients were recruited at the Institute of Gastroenterology, Nutrition and Liver Diseases at Schneider Children’s Medical Center of Israel. The study was approved by the hospital institutional review board (approval no. 0693-22) and written informed consent was obtained from parents or other legal guardian for all study participants. Anti-tTG values (Bioplex 2200, Bio-Rad Laboratories, CA) were obtained from the electronic health record for all participants, except participant P038, who had IgA deficiency. Endoscopies were performed as part of routine clinical care, in patients with suspected CeD (typically based on positive TTG serology with or without additional symptoms) or in patients referred for abdominal pain or nausea. All patients in the CeD group were eventually diagnosed with CeD, based on typical histologic findings, while subjects in the control group had a normal macroscopic and histologic evaluation of the duodenum. Duodenal biopsies specimens were immediately snap-frozen in liquid nitrogen and stored at −80 °C until further processing. Samples lacking histological confirmation of inflammation from the corresponding intestinal segment were excluded from the study. For preparation of FFPE blocks, frozen samples were thawed and fixed in 4% formaldehyde for 24 h, followed by transfer to the Histology Unit at the Weizmann Institute of Science for paraffin embedding.

### VisiumHD

VisiumHD dataset was assembled from 8 CeD and 4 control biopsies. Sections (4 µm thick) from FFPE tissue blocks were mounted onto VisiumHD–compatible slides and processed for hematoxylin and eosin (H&E) staining according to the VisiumHD FFPE Tissue Preparation Handbook (CG000684). Brightfield whole-slide images were acquired using a Leica DMi8 widefield inverted microscope equipped with a Leica DFC7000T color camera and an HC PL APO 20×/0.80 dry objective (506530; Leica Microsystems CMS GmbH). A Tile Scan acquisition was used to cover the full tissue area, and tiles were merged into a single image. Following imaging, tissue sections were processed for spatial transcriptomics using the VisiumHD Spatial Gene Expression Reagent Kits (CG000685) and CytAssist system (v2.1.0.14) at 37 °C for 30 min, in accordance with the manufacturer’s instructions. Generated libraries were loaded onto individual lanes of an Illumina NovaSeq X platform using a 1.5B flow cell at a final loading concentration of 150 pM.

### Single-nuclei isolation from tissue samples

The snRNAseq dataset was assembled from 6 CeD and 5 control biopsies. Two snap-frozen biopsy samples (P116 and P123) were processed according to the Tissue Fixation & Dissociation for GEM-X Flex Gene Expression protocol (CG000783). The remaining samples, comprising 100–200 µm of tissue obtained as serial 50 µm sections from FFPE blocks, were processed using the Sample Preparation from FFPE Tissue Sections for GEM-X Flex Gene Expression protocol (CG000784). Isolated single nuclei were subsequently processed following the GEM-X Flex Gene Expression Reagent Kits for Multiplexed Samples protocol (CG000787), in accordance with the manufacturer’s instructions. Prepared libraries were loaded onto Illumina NovaSeq X platform using a 1.5B flow cell at a final loading concentration of 180 pM with a 10% PhiX spike-in.

### Computational analysis

VisiumHD spatial transcriptomics sequencing data were processed using 10x Genomics Space Ranger software v3.1.2, while GEM-X Flex snRNAseq gene expression sequencing data were processed using Cell Ranger v9.0.0, in accordance with the manufacturer’s recommended workflows. For each cell in the snRNAseq and pixel in the VisiumHD datasets, normalized counts are the counts of each gene divided by the total number of UMIs for this cell/pixel. Next, the data was multiplied by 10,000 and logged using log1p function in python. Highly variable genes had a minimum expression threshold of 1e-5.

### Processing of VisiumHD data

#### Segmentation and classification of H&E images

Manual segmentation of hematoxylin and eosin (H&E) - stained images was performed using QuPath v0.5.1, restricting the analysis to regions with relatively preserved crypt–villus axis morphology. The muscularis mucosa/lamina propria/crypt (MM/LP/crypt) segment was defined as the region extending from the base of the villus segment to the muscularis mucosa. The crypt–villus border was defined as a line connecting the most basal luminal (non-tissue) pixels between adjacent villi or, in the case of blunted villi, as a line drawn immediately above the epithelial cell layer. In addition, Brunners’ glands and immune follicles were manually segmented to facilitate their exclusion from analyses such as zonal reconstructions. Spatial distances were calculated within predefined manually annotated “macrozones” to preserve the directional organization of tissue morphology. Classification of epithelial and non-epithelial pixels in H&E-stained sections was performed using the pixel classifier in QuPath software v0.5.1^39^. Random Trees pixel classifiers at Moderate resolution were trained using all available multiscale image features provided by the software (scales: 1,2,4,8) and subsequently applied to the annotated tissue regions.

#### Annotation and clustering of single pixel gene expression

For VisiumHD spatial transcriptomics data, only 8*8µm pixels containing a minimum of 100 unique molecular identifiers (UMIs) were retained for downstream analysis. Annotation was restricted to pixels classified by the pixel classifier and located within manually annotated villus, crypt, muscularis mucosa, or lamina propria regions. Samples with fewer than 1,000 pixels remaining after this initial filtering step were excluded from further analysis. Following normalization, highly variable genes were identified, expression values were scaled, and dimensionality reduction was performed using principal component analysis (PCA). A neighbor graph was constructed using n_neighbors = 10 and n_PCs = 40, followed by clustering with the Leiden algorithm at a resolution of 1.5. Cluster identities were manually annotated based on known markers.

#### Processing 10x Flex snRNAseq data

Background signal was removed on a per-sample basis by calculating the mean expression profile of low-UMI cells (100–500 UMIs) and subtracting this background from the corresponding sample counts. Following background subtraction, negative values were set to zero. Quality control metrics, including the fraction of mitochondrial transcripts, were computed using sc.pp.calculate_qc_metrics. Cells with 500–10,000 UMIs and a mitochondrial transcript fraction ≤10% were retained for downstream analysis.

Initial dimensionality reduction and clustering were performed using 30 principal components (n_PCs = 30) and a neighbor graph with n_neighbors = 10, followed by Leiden clustering at a resolution of 0.2. Cell lineages, including epithelial, stromal, and immune populations, were identified and subsequently subclustered to resolve finer cell types. During this process, clusters suspected of representing doublets were excluded, as were cells exhibiting aberrant expression of markers from other lineages, indicative of potential cross-contamination. Reclustering was performed for each lineage after removal of contaminated clusters. Final UMAP embeddings were generated using 15 principal components and 10 neighbors. GSEA analysis was performed using GSEApy ^40^.

#### Identification of Enterocyte Zonation Marker Genes

To identify marker genes associated with enterocyte zonation, enterocyte genes were first determined from the snRNAseq dataset using the rank_genes_groups function. Genes were retained based on a minimum log fold change of 2 and an adjusted p-value ≤0.01. This gene set was intersected with genes exhibiting higher expression in the villus compared to the crypt (villus-to-crypt expression ratio >1) and a mean expression level ≥1×10⁻⁴ in either crypt or villus epithelial pixels. Genes were subsequently ranked according to their center of mass (COM), computed as the gene-expression weighted distance bin along the crypt–villus axis^5^. The top and bottom 40 genes were selected as markers of villus tip and villus bottom enterocytes, respectively. Mid-villus marker genes were defined as genes exhibiting minimal expression of at least 1×10⁻⁴ in the mid-villus(250 µm) bin and preferential expression in this region, with a mid-villus–to–non–mid-villus expression ratio exceeding 1.

#### Zonation reconstruction and stromal cluster analysis

Zonation analysis was performed using only pixels annotated as villus or muscularis mucosae (MM)/lamina propria (LP)/crypt, while pixels corresponding to Brunner’s glands and immune structures were excluded. Samples were included only if they exhibited a minimum median of 20 UMIs per 8*8µm pixel. For each sample, distances were calculated relative to the crypt–villus border, with positive distances assigned to villus regions and negative distances to MM/LP/crypt regions. Distances were measured up to 800 µm and discretized into 100 µm bins.

For each distance bin, normalized gene expression was computed by summing pixel-level counts and dividing by the total counts within the bin. To account for variability in sample size, mean expression profiles for disease conditions (celiac disease and control) were computed using only distance bins present in at least 50% of samples. To identify stromal genes with extreme spatial localization, differentially expressed genes from stromal snRNAseq data were identified using the rank_genes_groups function in Scanpy. Genes with a normalized mean expression <1×10⁻⁵ were excluded. The remaining genes were ranked by their COM values based on the VisiumHD data. A curated list of WNT and BMP signaling molecules, excluding receptors and transcription factors, from Harnik et al.^5^ was intersected with genes falling within the top and bottom 10% of COM values. COM peak profiles of the summed expression of top and bottom gene sets were calculated on high quality enterocytes with at least 1,000 UMIs across all patients as well as on a per-patient basis. Stromal marker genes were identified based on a minimum mean expression threshold of 1×10⁻⁴ and a minimum fold change of 2 between the mean expression of a given stromal cell type and the maximal mean expression observed across all other cell types. Analyses in Figs. S2E, F were performed on means of all VisiumHD pixels.

#### HCR-FISH

Samples were stained according to the Molecular Instruments HCR GOLD protocol for FFPE tissues. In brief, FFPE sections were deparaffinized and rehydrated through a graded ethanol series (Xylene, Xylene, 100% Ethanol, 100% Ethanol, 95% Ethanol, 70% Ethanol, 50% Ethanol, PBS). Sections were permeabilized for 10 min with 0.5% TritonX-100 (Sigma Aldrich) followed by two washes with PBS. Tissues were incubated in HiFi HCR hybridization buffer for 10 min in a 37 °C incubator. Tissues were incubated with hybridization buffer mixed with probes (1:50) for 16 h in 37 °C. After the hybridization, tissues were washed with HiFi HCR wash 15 min in 37 °C for 4 times. The samples were incubated with HCR amplification buffer (AB) for 30 min at room temperature. HCR-hairpins (H1 and H2 hairpins of X1-546, B5-647) were heated for 90 s at 95 °C and cooled in dark for 30 min. The samples were incubated with AB with 6 pmol of each hairpin for 60 min in the dark, and then rinsed twice in GOLD HCR wash buffer for 15 min. The samples were incubated with GOLD HCR wash buffer with DAPI (50 ng ml−1, Sigma-Aldrich, D9542) and a488 wheat germ agglutinin (WGA, 1:1000, ThermoFisher scientific W11261) for 5 min. Additional wash with GOLD HCR wash buffer was performed, and slides were mounted with ProLong Gold Antifade Mountant (ThermoFisher scientific, P36934). Images were acquired using 90× magnification on a Nikon-Ti-E inverted fluorescence microscope using the NIS element software AR 5.02.03.

## Supplementary Figures

**Supplementary Figure 1.**
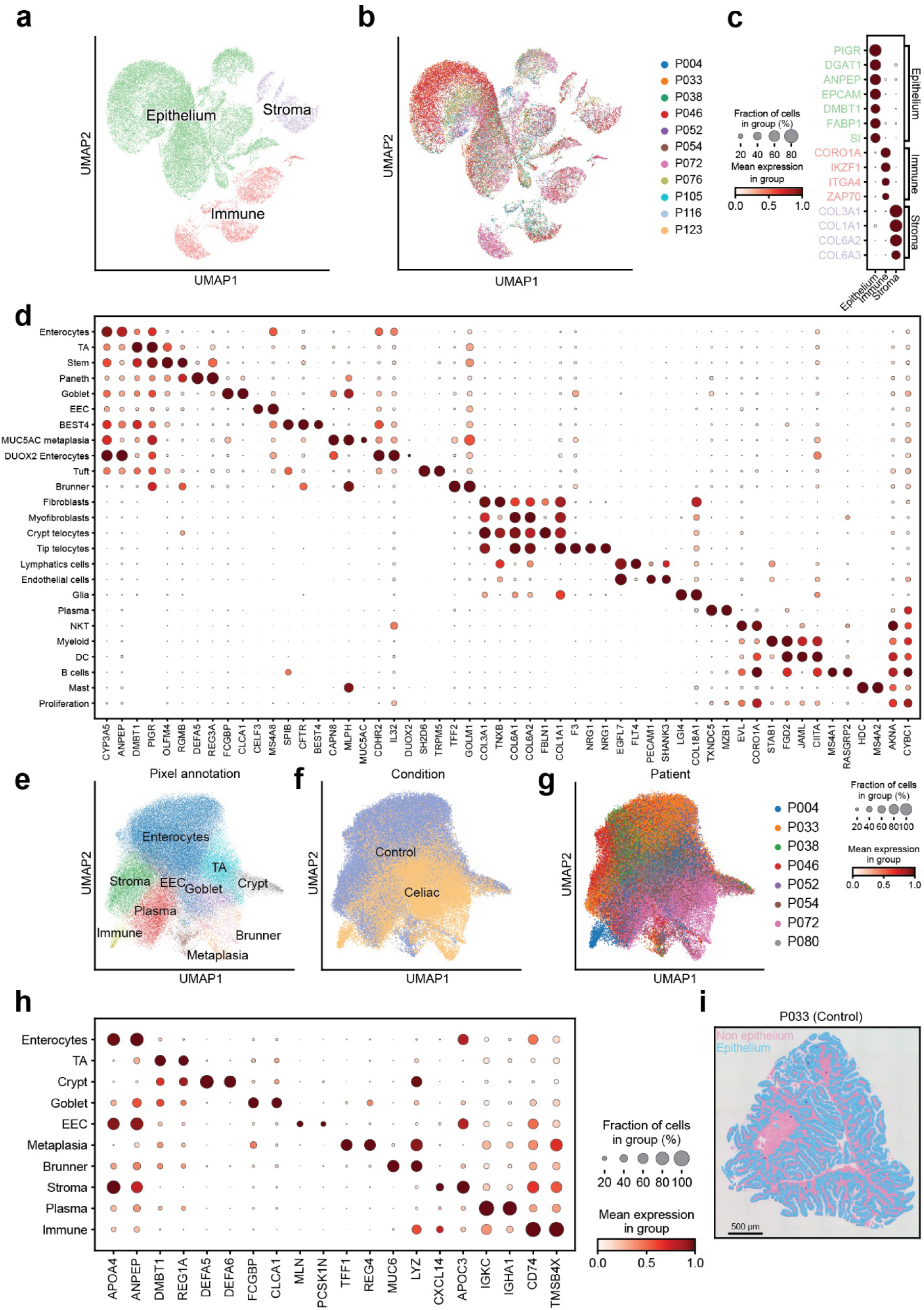
Single-cell and spatial transcriptomic characterization of pediatric duodenal tissue in control and in CeD **a.** UMAP projection of snRNAseq data showing broad cell type annotations. **b.** UMAP projection of snRNAseq data colored by patient. **c.** Dot plot displaying marker gene expression for each broad cell type identified in the snRNAseq dataset. **d.** Dot plot showing marker genes defining fine-grained cell types in the snRNAseq data. **e.** UMAP projection of VisiumHD spatial transcriptomics data showing pixel-type annotations. **f.** UMAP projection of VisiumHD data colored by disease condition. **g.** UMAP projection of VisiumHD data colored by patient. **h.** Dot plot showing marker gene expression across pixel types in VisiumHD data. **i.** Representative example of pixel classification, illustrating epithelial (light blue) and non-epithelial (pink) pixels. In c,d,h – Expression data were standardized using min–max scaling (0–1), performed separately for each gene.

**Supplementary Figure 2.**
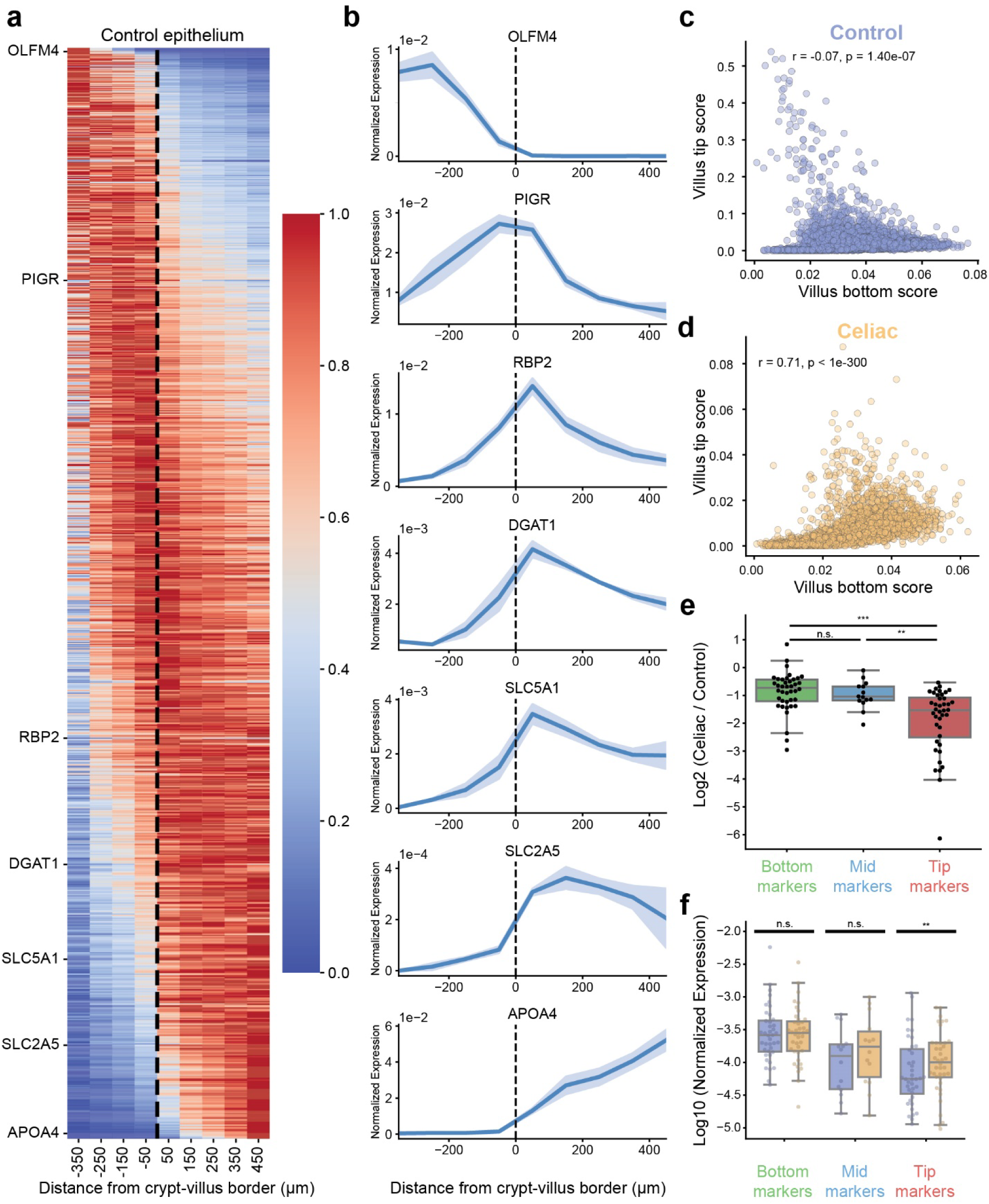
Spatial reconstruction of epithelial zonation along the crypt-villus axis **a.** Zonation across control epithelial samples (*n* = 4). Each column represents a 100 µm spatial bin and each row represents a gene. Gene expression values were smoothed, max-normalized across zones, and sorted by COM. **b.** Representative expression profiles of zonated genes shown in a. **c.** Scatter plot of control enterocytes showing a negative correlation between bottom and tip zonal scores (Methods). **d.** Scatter plot of CeD enterocytes showing a positive correlation between bottom and tip zonal scores. **e.** Pseudobulk analysis of VisiumHD epithelial 8*8µm pixels showing log₂ fold changes in gene expression between CeD and control samples. **f.** Pseudobulk expression of bottom, mid, and tip enterocyte markers in crypt epithelial pixels, demonstrating increased expression of tip markers in CeD. In e,f - Statistical significance was assessed using the Wilcoxon rank-sum test; ** and *** denote *p* < 0.01 and *p* < 0.001, respectively. Box plots show the median as the center line, boxes span the 25th–75th percentiles, whiskers extend up to 1.5 IQR.

**Supplementary Figure 3.**
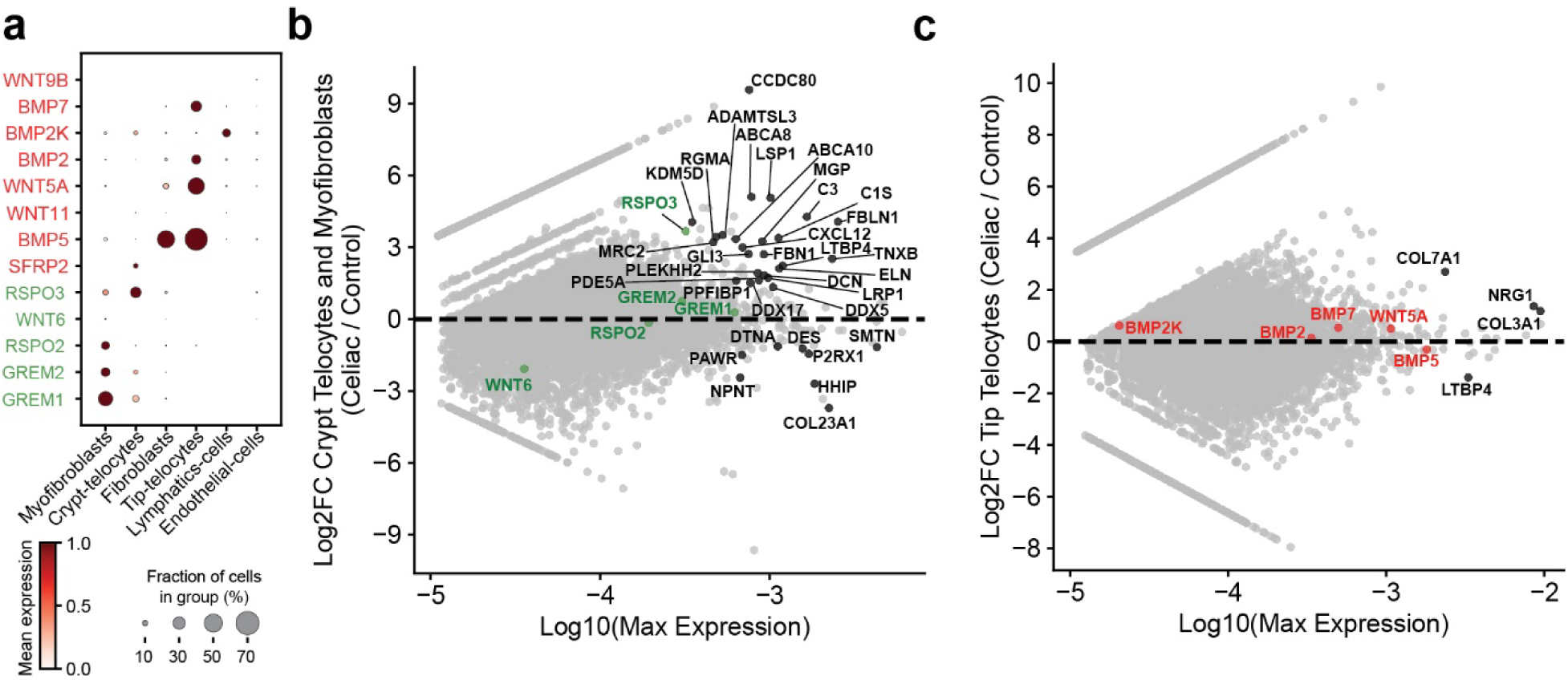
Stromal morphogens expression along the crypt-villus axis **a.** Dot plot showing expression of tip- and crypt bottom-associated morphogens across stromal cell populations in the snRNAseq data set. Expression data were standardized using min–max scaling (0–1), performed separately for each gene. **b.** MA plot of crypt-telocytes and myofibroblasts showing expression of bottom morphogens, indicating the absence of differential morphogen expression between CeD and control samples. **c.** MA plot of tip-telocytes showing expression of tip morphogens, indicating the absence of differential morphogen expression between celiac disease and control samples. Black dots in b-c show significantly differentially expressed genes, none of the crypt (green) and tip (red) morphogens were significantly differentially expressed in CeD.

**Supplementary Figure 4.**
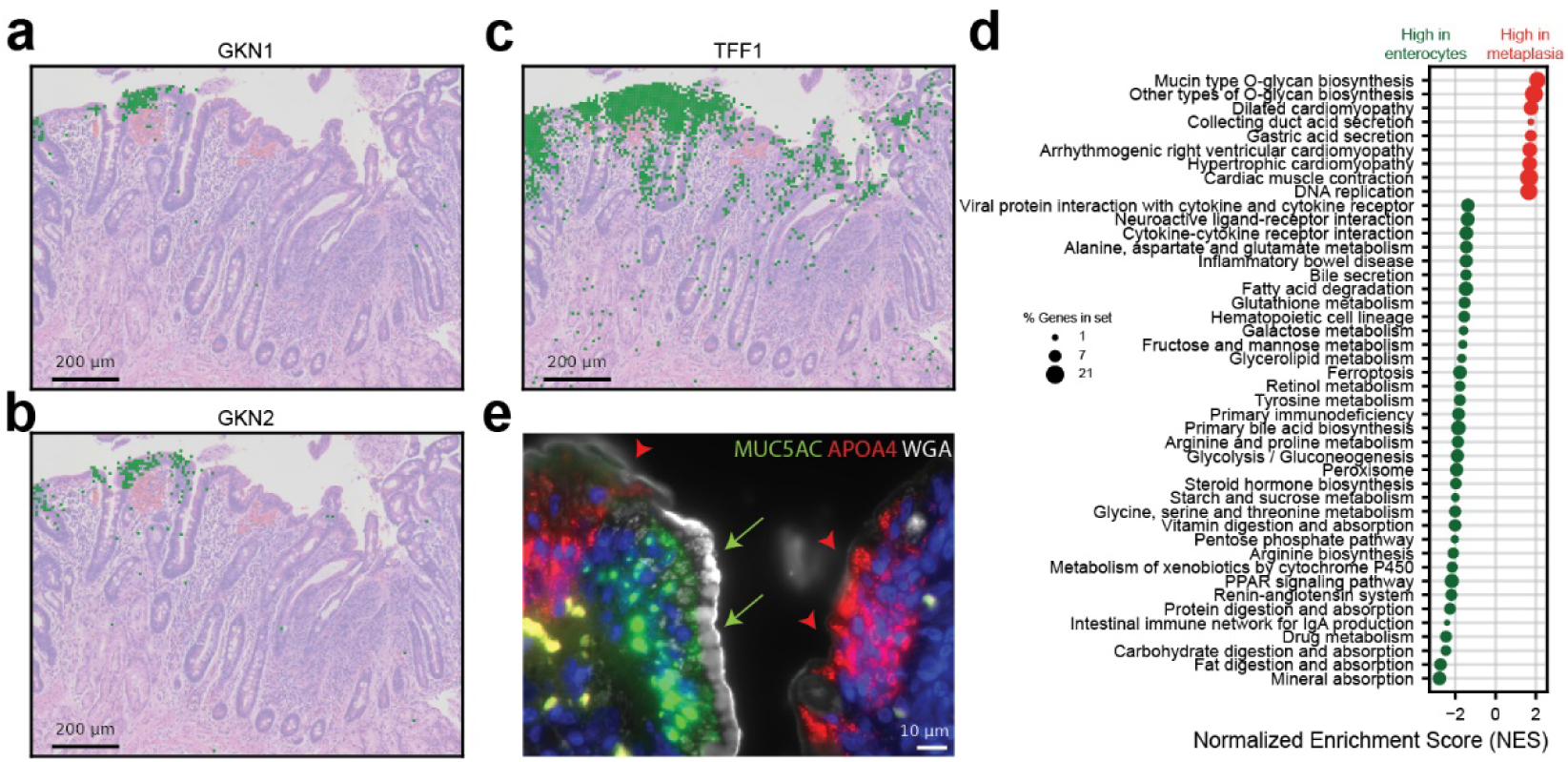
Spatial and transcriptional characterization of gastric metaplasia in CeD **a-c.** Spatial transcriptomics image from a CeD patient highlighting positive pixels positive for gastric metaplasia associated genes, *GKN1, GKN2, TFF1*. **d.** GSEA showing transcriptional programs in metaplastic cells compared with CeD enterocytes. Shown are gene sets with q-value<0.2 **e.** Hybridization Chain Reaction Fluorescence in situ hybridization (HCR-FISH) image showing metaplastic cells (green, *MUC5AC* positive) and enterocytes (red, *APOA4* positive), and wheat germ agglutinin (WGA) staining (white) demonstrating higher mucin content in the metaplastic cells (green arrows), compared with the adjacent enterocytes (red arrowheads).

## Supplementary tables

**Supplementary Table 1.** Duodenal zonation patterns in control and pediatric celiac samples. Shown are the means and standard errors of the means (se) across patients for epithelium, non-epithelium and all pixels. Com_norm – center of mass normalized between 0 and 1. **Related to Figure 2**.

**Supplementary Table 2.** Gene expression markers for the villus bottom, mid and tip. **Related to Figure 2**.

